# Detection of transient neurotransmitter response using personalized neural networks

**DOI:** 10.1101/2019.12.19.883017

**Authors:** Ivan S. Klyuzhin, Connor W. J. Bevington, Ju-Chieh (Kevin) Cheng, Vesna Sossi

**Affiliations:** Department of Medicine University of British Columbia Vancouver, BC, V6T 1Z3, Canada; Department of Physics and Astronomy University of British Columbia Vancouver, BC, V6T 1Z4, Canada

**Keywords:** Neurotransmitter response, dopamine release, lp-ntPET, neural network, brain imaging, PET

## Abstract

Measurement of stimulus-induced dopamine release and other types of transient neurotransmitter response (TNR) from dynamic PET images typically suffers from limited detection sensitivity and high false positive rates. Measurement of TNR of a voxel-level can be particularly problematic due to high image noise. In this work, we perform voxel-level TNR detection using artificial neural networks (ANN) and compare their performance to previously used standard statistical tests. Different ANN architectures were trained and tested using simulated and real human PET imaging data, obtained with the tracer [^11^C]raclopride (a D2 receptor antagonist). A distinguishing feature of our approach is the use of “personalized” ANNs that are designed to operate on the image from a specific subject and scan. Training of personalized ANNs was performed using simulated images that have been matched with the acquired image in terms of the signal and noise.

In our tests of TNR detection performance, the F-test of the linear parametric neurotransmitter PET (lp-ntPET) model fit residuals was used as the reference method. For a moderate TNR magnitude, the areas under the receiver operating characteristic curves in simulated tests were 0.64 for the F-test and 0.77–0.79 for the best ANNs. At a fixed false positive rate of 0.01, the true positive rates were 0.6 for the F-test and 0.8–0.9 for the ANNs. When applied to a real image, the ANNs identified a TNR cluster missed by the F-test. The newly found cluster was verified to contain TNR by direct lp-ntPET model fitting. These results demonstrate that personalized ANNs may offer a greater detection sensitivity of dopamine release and other types of TNR compared to previously used methods.

## 1 Introduction

Neurological and psychiatric disorders are often associated with alterations in multiple neurotransmitter systems. For example, in Parkinson’s disease, degeneration of the dopaminergic system is known to be largely responsible for the motor symptoms, while changes in the serotonergic [1] and cholinergic [2] systems are mostly related to mood and cognition. In subjects with obsessive-compulsive disorder, the binding of serotonin transporter in the insular cortex was found to be decreased compared to healthy controls [3]. In populations prone to risk-taking and other compulsive behaviors, impaired reward processing has been found to be associated with abnormal stimulus-induced dopamine release [4]. Probing the function of the neurotransmitter systems in these and other disorders can provide important insights into disease etiology and mechanisms.

Stimulus-induced changes in the synaptic neurotransmitter levels can be estimated using dynamic positron emission tomography (PET) imaging with receptor radioligands. The most widely used approach is to compute the difference in the non-displaceable binding potential (ΔBP_ND_) [5] of a radioligand between a baseline and a post-stimulus scan. A limitation of this method is that it assumes that the neurotransmitter system is in a steady state during the scans; thus, when there is a transient change in neurotransmitter concentration, also termed transient neurotransmitter response (TNR), factors such as the magnitude and duration of the TNR cannot be determined [6]. To address this shortcoming, a linear parametric neurotransmitter PET model (lp-ntPET) has been developed [7,8]: by fitting the lp-ntPET model to the regional time-activity curves (TACs), one can determine the precise time course of TNR from a single scan.

Applying lp-ntPET on a voxel level can reveal a detailed spatial distribution of TNR [8]. However, such application becomes problematic in high-resolution PET due to a typically high image noise, which originates from a small number of acquired counts per voxel. The problem is exacerbated by the requirement to acquire dynamic PET data at a relatively high temporal sampling rate (1–3 minutes per frame), further reducing the number of counts. Fitting lp-ntPET on such data will almost always produce non-zero TNR, regardless of the truth [9]. To identify voxels that are more likely to contain true TNR, the F-test is performed between two model fits: a baseline model that does not account for TNR, typically the simplified reference tissue model (SRTM) [10], and lp-ntPET. However, the use of the F-test in this application may suffer from several limitations; first, it does not exploit any possibly useful prior assumptions about the TAC shape. Secondly, F-test assumes normally-distributed samples, while the noise in iteratively-reconstructed PET images is known to be closer to a gamma distribution [11]. Finally, it does not fully account for the different noise levels present in different frames. To reduce the impact of noise, previous work has explored direct reconstruction of lp-ntPET parametric images from acquired PET data [12]: while this method helped to improve the accuracy and precision of the kinetic parameter estimates, the fraction of false positive activations remained relatively high.

We have previously demonstrated that relatively simple artificial neural networks (ANNs) outperform conventional methods of dynamic PET denoising [13]. Based on this finding, in this work we propose to use ANNs for the task of voxel-level TNR detection in reconstructed images, as a substitute for the F-test. Unlike the F-test, ANNs can efficiently encapsulate both the signal and noise properties of the training data, and the noise is not required to follow any particular statistical distribution. On the other hand, the ANN output was found to be rather sensitive to the change in the input distribution, i.e. TAC shapes not present in the training set were not processed correctly.

The novelty of our present work is that, for the first time, we apply various types of ANNs to detect TNR in simulated and real human data. Further, we propose that the high specificity of ANNs to the training set can be used as an advantage: in order to maximize the TNR detection sensitivity in noisy voxel TACs, we train ANNs to only operate on a single dynamic data set obtained for a particular subject and scan. We term such ANNs “personalized” (pANN), although it is understood that such networks are also scan-specific in addition to being subject-specific. Finally, we test whether ANNs trained entirely on simulated data (that have been matched with real data) can generalize to real data without significant errors and abnormalities.

To construct a pANN for TNR detection in a particular scan, a reference-region TAC and governing kinetic parameters are first determined from the acquired scan, and then used to generate well-matched simulated data for pANN training. As long as the employed ANN architecture is relatively small, the network can be readily trained using the simulated data and applied to the acquired dynamic image within minutes, without imposing significant limitations on the image processing and analysis throughput. Thus, similarly to our previous denoising work [13], we focused on testing relatively small ANNs with standard architectures: a fully-connected neural net, a convolutional neural net, as well as two architectures based on denoising autoencoders where the TNR signals were treated as anomalies in baseline TACs. All networks were trained to output the non-calibrated confidence of TNR presence in a given voxel’s TAC. In order to objectively compare the pANNs to the F-test, dynamic test images with known ground truth were generated by simulating tracer kinetics and PET data projection, according to an established methodology [14, 15]. The simulated test data have been matched with acquired data from a scan with [^11^C]raclopride (RAC, a dopamine D2 receptor antagonist) where transient dopamine release was induced mid-scan by means of a gambling task. Several performance metrics were evaluated and compared between the F-test and pANNs: 1) areas under the receiver operating characteristic (ROC) curves (AUC) for different BP_ND_ values, TNR magnitudes, and TNR cluster sizes; 2) TNR detection sensitivity, fraction of detected TNR clusters, and Dice coefficients at fixed false positive (FP) rates; 3) the size distributions of FP clusters. Finally, we applied the pANNs and the F-test to detect TNR in the acquired RAC scan, and compared the results.

## 2 Methods

### 2.1 Method overview

The pipeline for training and applying a pANN for TNR detection is illustrated in Fig. 1a. A reconstructed image from the RAC scan was denoised using a generic method, and a tissue-derived input function (reference TAC) was measured from the image. Using SRTM2 [16], the distributions of kinetic parameters in the non-specific and specific tracer binding regions were determined. Next, lp-ntPET model was used to generate a noise-free dynamic (4D) image from the reference TAC and randomly-generated volumetric map of kinetic parameter distributions (Fig. 1b). For voxels containing simulated TNR, the lp-ntPET parameters were set according to the scan protocol and stimulus timing. The dimensions, temporal sampling, and voxel size of the generated image matched those of the acquired image.

**Figure 1:**
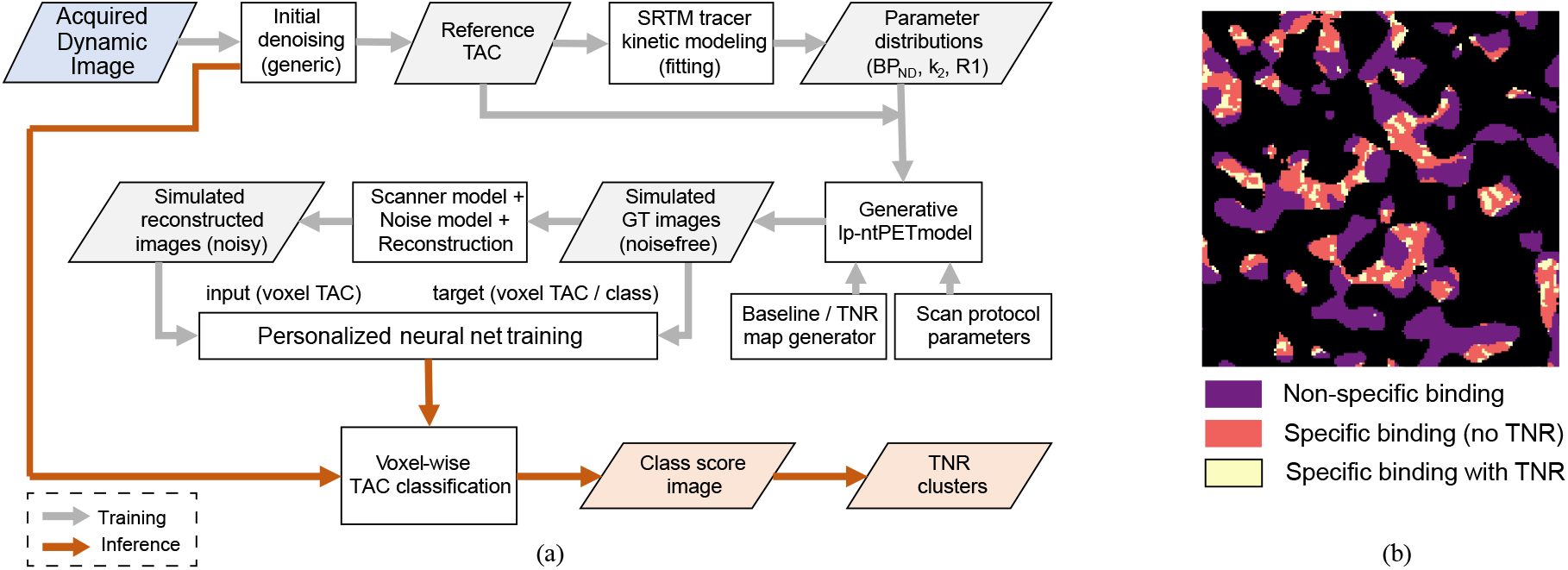
(a) The pipeline for training a pANN for voxel-level TNR detection. (b) A randomly-generated volumetric map (single plane is shown) with regions denoting non-specific and specific tracer binding, as well as regions of specific binding with TNR.

The noise-free image was forward-projected using the system model of the PET scanner, Poisson noise was added to the projection data, and the data were reconstructed and post-processed with the same methods as the acquired data to obtain a noisy dynamic image. Voxel TACs from the noisy image were used in the pANN training as inputs, and voxel classes (baseline/TNR) were used as targets. We then applied the pANN to the acquired dynamic image (or simulated test images) to obtain a parametric class score image. The class score image was thresholded to obtain the TNR cluster map. Below we provide a detailed description of these steps.

### 2.2 Image acquisition and post-processing

The healthy subject RAC scan was performed on the high resolution research tomograph (HRRT, Siemens) after a bolus injection of 20 mCi of RAC. The scan duration was 75 min; at 36 minutes into the scan, the subject was presented with a stimulus (Vancouver gambling task [17]) to induce transient dopamine release in the striatum. The scan duration and stimulus timing were chosen according to a previously published optimized protocol [9]. The acquired data were histogrammed into 30 temporal frames (frame durations 5× 1 min, 5×2 min, 20×3 min).

The acquired data were reconstructed using the ordered-subset expectation maximization algorithm with incorporated highly-constrained backprojection operator (HYPR-OSEM) [18] (16 subsets, 6 iterations); the HYPR kernel size was (5.0 mm)^3^. The image dimensions were 256×256×207 voxels, with an isotropic voxel size of (1.22 mm)^3^. After image reconstruction, HYPR-based denoising [19] with a (5.0 mm)^3^ kernel size was applied to further reduce noise. The subject was also scanned on a Philips Achieva 3T scanner to provide anatomical reference. The acquired T1-weighted image was segmented using Freesurfer 6.0 [20] to obtain masks of the cerebellum and striatum.

### 2.3 Kinetic models for dynamic PET Data

As mentioned above, we used two models to fit TACs from the acquired scan and to simulate RAC kinetics. SRTM was used to analyze the baseline state:

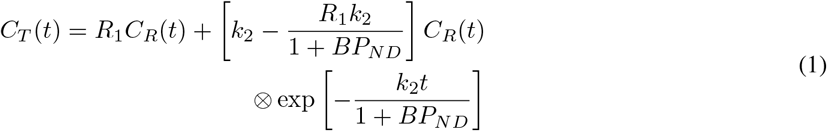

where *C_T_*(*t*) is the measured (fitted) TAC, *t* is the frame time, k_2_ is the tissue-to-plasma rate constant, R_1_ is the relative delivery in the target region compared to the reference region, *C_R_*(*t*) is the tissue-derived input function (reference region TAC); ⊗ denotes the temporal convolution. From this equation, a noise-free TAC with no TNR can be generated given the input function and the triplet of values BP_ND_, k_2_, R_1_.

To model TACs with TNR, we used the lp-ntPET model [7]:

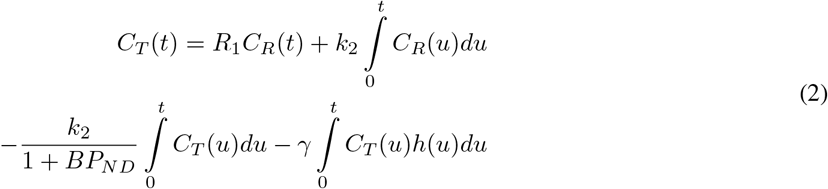

where the last term accounts for time-varying TNR, and *γ* is related to the magnitude of TNR. *h*(*t*) is the time-course of TNR modeled as the function

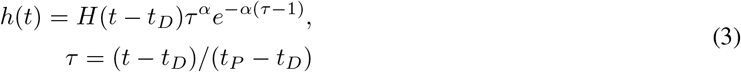

where *H* is the Heaviside step function, *t_D_* and *t_P_* are the start and peak times of TNR, respectively; the parameter *α* determines the sharpness of the response. The fitted parameters include BP_ND_, k_2_, R_1_, *γ*, *t_D_, t_P_*, and *α*.

### 2.4 Training image generation

The cerebellar white matter, defined by a mask produced by Freesurfer, was used as reference region to determine the tissue input TAC *C_R_*(*t*) from the acquired RAC scan. SRTM2 [16] was applied on the voxel level, and the joint distributions of BP_ND_, k_2_ and R_1_ in the non-specific (cerebellum) and specific (striatum) binding regions were evaluated. To avoid biased parameter values due to the presence of TNR, the SRTM2 was fit only to the TACs limited to the first 36 minutes, i.e. prior to the stimulus. By comparing full-length lp-ntPET fits and 36-minute SRTM2 fits, it was verified that using the shortened TACs for SRTM2 fitting did not introduce bias in the kinetic parameters. The 1st and 99th percentiles of the measured BP_ND_ distribution were (−0.45, 0.89) in the cerebellum, and (0.78, 5.72) in the striatum. The corresponding percentiles for k_2_ were (0.0015, 0.009) in the cerebellum, and (0.0022, 0.0065) in the striatum. The ratio of R_1_/k_2_ for the subject was 260.0 for all voxels; thus we set *R*_1_ = 260 × *k*_2_.

To generate simulated dynamic images for pANN training, we started with a random volumetric map, which labeled 3D regions as corresponding to non-specific, specific and zero tracer binding (i.e. cold regions) (Fig. 1b). Since the tested ANNs had one-dimensional input (single-voxel TAC, similar to the F-test), there was no need to accurately model anatomical structures in the brain. However, the general shapes and dimensions of the random regions were similar to anatomical structures in typical acquired RAC images, as has been previously described in detail [13]. This helped equalize the impacts of resolution blurring and neighboring-voxel correlation in the simulated and real images.

Voxels in the generated regions were assigned triplets of (BP_ND_, k_2_ and R_1_) values randomly drawn from a joint uniform distribution, where the distribution boundary was determined from the acquired image. Following the value assignment, an anisotropic diffusion filter [21] was applied, which produced smoothly varying local gradients of the kinetic parameters (magnitudes and directions) within the specific and non-specific binding regions, while preserving high-contrast edges between the regions.

The specific binding regions were set to contain clusters of TNR, ranging in size from 100 to approximately 500 voxels. Each cluster had unique randomly assigned values of *γ*, *t_D_, t_p_* and *α*, set in accordance with the stimulus timing [8,9,22]:

- *γ* ranged from 0.1 × 10^−3^ to 1.0 × 10^−3^ s^−1^.
- The range of *t_D_* (TNR start time) was 36 ± 5 min.
- The range of *t_P_* (TNR peak time) was from *t_D_* +5 to *t_D_*+15 min.
- *α* was randomly chosen between 0.25, 1.0 and 4.0.

Based on the assumption that the profiles of TNR in nearby voxels should be similar, the values of these parameters were set to be spatially constant within each cluster.

From the generated parametric images, we simulated noise-free TACs for each voxel using (2) and the reference TAC from the acquired scan. For the voxels with no TNR, the parameter *γ* was set to zero. The simulated TACs were resampled into 30 time points, to yield dynamic images with frame number and durations consistent with those in the acquired RAC image. Likewise, the voxel size was set to (1.22 mm)^3^, equal to the HRRT image voxel size.

To simulate intrinsic resolution blurring of the HRRT, each frame of the simulated noise-free image was smoothed with an isotropic 2.5 mm full-width at half-maximum (FWHM) Gaussian filter [23]. Each frame was then analytically forward-projected using the HRRT system matrix, and the effects of subject attenuation and detector normalization were included in the sinogram space. Random coincidences or scatter were not included in the simulation, since with real HRRT images these effects are taken into account during the reconstruction. Poisson noise was added to the forward-projected 3D sinogram data, which were reconstructed and post-processed similarly to the acquired image (Fig. 2a).

**Figure 2:**
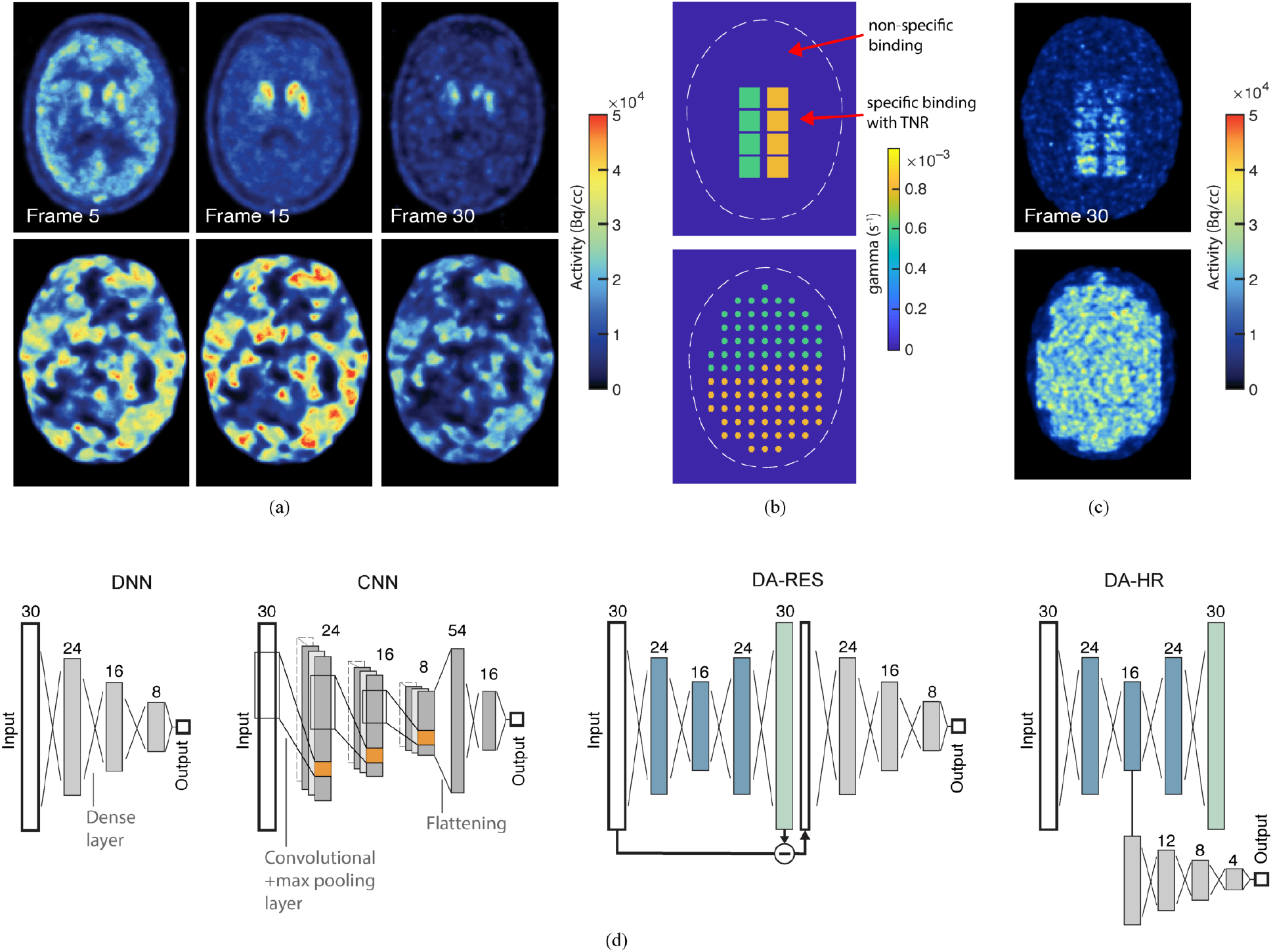
(a) Examples of frames from the acquired RAC image (top) and simulated training images (bottom). Single planes are shown. (b) Examples of parametric test images showing the regional distribution of *γ* values. Uniform regions and TNR clusters with *γ* = 0.2 × 10^−3^ and 0.4× 10^−3^ were included in planes that are not shown. (c) Examples of frames from the simulated dynamic test images. The shown planes correspond to those in panel (b). (d) Architectures of the tested ANNs for TNR detection. The autoencoder layers are shown in blue. The number of nodes per layer is shown above the layer blocks.

### 2.5 Test image generation

Two parametric test images were used, containing two distinct test patterns: large regions with uniform TNR, and spherical TNR clusters (Fig. 2b). Real TNR patterns are expected to fall in-between these two cases.

In the first image, the values of *γ* were set to be different in different uniform regions, equal to 0.2× 10^−3^, 0.4× 10^−3^, 0.6× 10^−3^ and 0.8× 10^−3^. The values of *t_D_* and *t_P_* were set in all regions to 36 and 46 minutes, respectively, and *α* was set to 1.0.

In the second image, TNR clusters of various sizes were placed in a baseline “background” with BP_ND_ set to 4.0. The cluster diameters were 3 voxels (3.6 mm), 5 voxels (6.0 mm) and 7 voxels (8.4 mm) (the plane shown in Fig. 2b only contained the 5-voxel sources). The values of *γ* in the clusters were similarly set to 0.2× 10^−3^, 0.4× 10^−3^, 0.6× 10^−3^ and 0.8× 10^−3^; the other TNR parameters were similar to those in the uniform regions. The procedure to generate noisy dynamic images for testing was similar to that for the training image; examples of individual frames are shown in Fig. 2c. The voxel TACs were generated using the measured reference TAC.

Initial experiments have shown that using a (5.0 mm)^3^ HYPR reconstruction/post-processing kernel produced strong ceiling and floor effects when measuring the classification performance in the uniform and cluster TNR regions, respectively. Therefore, with simulated test images, the HYPR reconstruction and post-processing kernel size was reduced to (2.5 mm)^3^; the ANN training data were adjusted accordingly.

### 2.6 Neural net architectures

Diagrams of the four tested neural net architectures are plotted in Fig. 2d. The first architecture was a traditional dense neural net (DNN) with three hidden layers. The number of nodes in each layer was 30 (input), 24, 16, 8, 1 (output). All layers had rectified linear unit (ReLU) activations, except the output, which had sigmoid activation (ranging from 0 to 1).

The second architecture was a one-dimensional convolutional neural net (CNN), with three convolutional layers followed by a dense layer and a regression layer with sigmoid activation. The number of neurons (or filters) in each layer was the same as the number of nodes in the DNN; a max pooling layer with a downscale factor of 2 followed each convolutional layer. The span of convolution windows was set to 1/2 of that layer’s input size (i.e. for the first layer the span was equal to 15).

The third architecture was based on a denoising autoencoder, with an auxiliary neural net operating on the input-output residuals (denoted DA-RES). This method was inspired by the use of autoencoders for anomaly detection, where the difference between the inputs and outputs (e.g. mean squared error) is often used as the anomaly score [24]. Within the context of our work, TNR signals can be treated as “anomalies” in normal baseline TACs. However, since we had simulated examples of such anomalies, instead of using an unsupervised anomaly metric we performed supervised training a small auxiliary neural net. The DA consisted of 5 fully-connected feed-forward layers, as illustrated. The number of nodes in each layer was 30 (input), 24, 16, 24, 30 (output), and all layers had ReLU activations.

In the fourth architecture, the auxiliary network was set to operate on the DA’s hidden representation (bottleneck) layer (denoted DA-HR). This design was inspired by a previous work on autoencoder-based anomaly detection, where a one-class support vector machine [25] or one-class neural net [26] were trained using the DA’s hidden data representation. With DA trained on baseline TACs, TNR TACs were expected to produce anomalous activations in the bottleneck nodes. Similarly to DA-RES, we perform supervised two-class training of the auxiliary network, which was intended to detect when the activation pattern was different from the normal distribution.

Using cross-validation, we evaluated the performance of DNNs/CNNs with different number of hidden/convolutional layers. The validation dataset was constructed by randomly picking 20% of the training data, and ten cross-validation folds were used. We found that there was very little benefit in using more than three hidden/convolutional layers in terms of the validation error. We also tested different convolution window spans in the CNNs equal to 1/3, 1/2 and 2/3 of the previous layer size, and found that the validation error was 1% lower on average in the latter two cases. Once the number of hidden layers was determined, we trained the networks for testing using the entire training data.

For DNN/CNN training, noisy baseline/TNR TACs were used as inputs, and the TAC classes (0 for baseline, 1 for TNR) were used as targets. The baseline TACs only included those corresponding to specific binding, since biologically no TNR can be expected in non-specific binding regions. The number of baseline and TNR training samples was 100000 each, and training was performed for 300 epochs.

The DA-based architectures were trained in two phases. In phase one, the DA network was trained using only baseline TACs: noisy TACs were used as inputs, and noise-free TACs were used as targets. Training was performed for 300 epochs using 100000 samples. In the second phase, the autoencoder weights and biases were fixed, and training of the auxiliary network was performed. In this phase a mixture and baseline and TNR TACs was used, and TAC classes were used as targets. The number of training samples was 100000 for each class, and training was performed for 200 epochs. We used the ADAM optimizer with the initial learning rate of 0.001; mean squared error was the optimized loss function.

### 2.7 Reference method

The following previously proposed [8] method to detect voxels with TNR was used here as the “reference”. The noisy TAC from a given voxel was fitted with the SRTM2 and lp-ntPET models, described by (1) and (2). The fit weights for different frames were set equal to the inverse of the decay correction coefficient for each frame [14]. The weighted residual sum of squares (wRSS) was computed for the two fits, and the F-score was computed using the equation

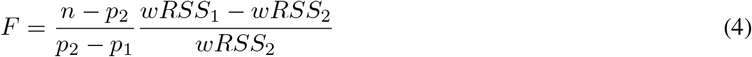

where *n* is the number of time samples in the TAC, the subscript designates the model (1 for SRTM2, 2 for lp-ntPET), and *p* is the number of parameters in the model. An empirically-determined threshold is applied to the F values to determine voxels with TNR. The F score can then be treated as the decision function of a classifier, and compared to other anomaly detection methods (e.g. by comparing the ROC AUC).

### 2.8 Method testing and comparison

Using the simulated test images, we computed several objective metrics that quantified the TNR detection performance, and compared them between DNN, CNN, DA-RES, DA-HR and the F-test. The ROC AUC was measured for the task of baseline/TNR classification of individual voxels. For voxels taken from the uniform baseline/TNR regions, the AUC was measured for different BP_ND_ and *γ* values; for voxels from the cluster regions, the AUC was measured for different *γ* values and cluster sizes.

For voxels in the uniform regions, we also measured the TNR detection sensitivity at fixed FP rates of 0.05 and 0.01. Likewise, for TNR clusters, the detected cluster fraction, cluster specificity, and Dice coefficient (using the detected and ground truth cluster shapes) were measured at FP rates of 0.05 and 0.01. These metrics were obtained and compared between the methods for different values of BP_ND_, *γ* and cluster sizes.

Using the uniform baseline regions, we measured the distribution of sizes of FP clusters, and estimated the cluster size threshold to reject 95% and 99% of FP clusters.

Finally, we applied the neural nets and the F-test to detect dopamine release in the acquired RAC image. Since in this case we don’t know the ground truth TNR distribution, we simply compare the TNR clusters detected by the neural nets to those detected by the F-test.

## 3 Results

### 3.1 ROC analysis

The ROC AUC values measured for voxels in the uniform TNR regions are plotted in Fig. 3a, for different values of BP_ND_ and *γ*. The graphs demonstrate that the methods DNN, CNN and DA-RES had consistently higher AUCs compared to the F-test; on the other hand, DA-HR performed similarly or worse than the F-test. For *γ* values of 0.4 × 10^−3^ and greater, the ANN methods offered only a small improvement over the F-test, likely due to approaching the maximum possible value of AUC. For *γ*=0.2× 10^−3^, the biggest improvement compared to the F-test was found for BP_ND_=4.

**Figure 3:**
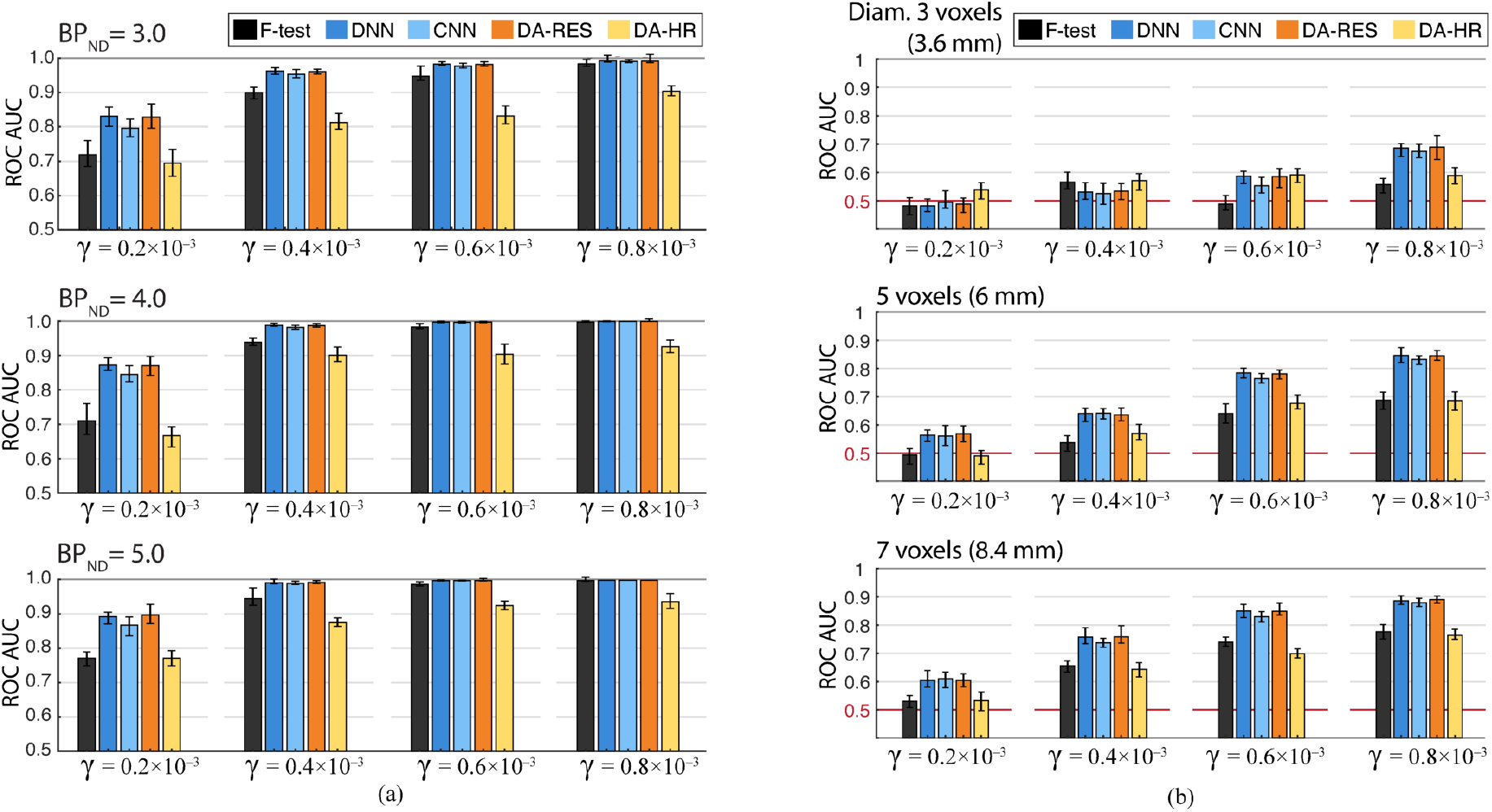
Comparison of baseline/TNR classification ROC AUC values between the tested methods. (a) AUC values measured in the uniform TNR regions. (b) AUC values measured in TNR clusters (BP_ND_=4).

The ROC AUC values measured for voxels in the TNR clusters are plotted in Fig. 3b. Here the AUC values were generally much lower compared to those for the uniform regions, likely due to the presence of the partial volume effect (i.e. TNR TACs from small clusters become mixed with baseline TACs from the surrounding background). The graphs demonstrate that DNN, CNN and DA-RES consistently and substantially outperform the F-test, with different cluster sizes and *γ* values; the AUC values of DNN and DA-RES are practically identical, while the values for CNN are marginally lower. DA-HR performed on average similarly to the F-test. None of the methods could effectively detect TNR in 3-voxel clusters with *γ* less than 0.8× 10^−3^.

Examples of the ROC curves for voxels in the uniform regions are plotted in Fig. 4a for *γ*=0.2×10^−3^ and 0.4×10^−3^. The curves demonstrate that there is indeed a significant improvement with DNN or CNN over the F-test (DA-RES curves were similar to those of DNN). The shapes of the ROC curves imply that all ANNs except DA-HR would offer a substantially better true positive rate, i.e. sensitivity, at practically any FP rate. Of particular interest are low FP rates of 0.05 and 0.01; in these regimes the F-test had better true positive to FP trade-off compared to DA-HR.

**Figure 4:**
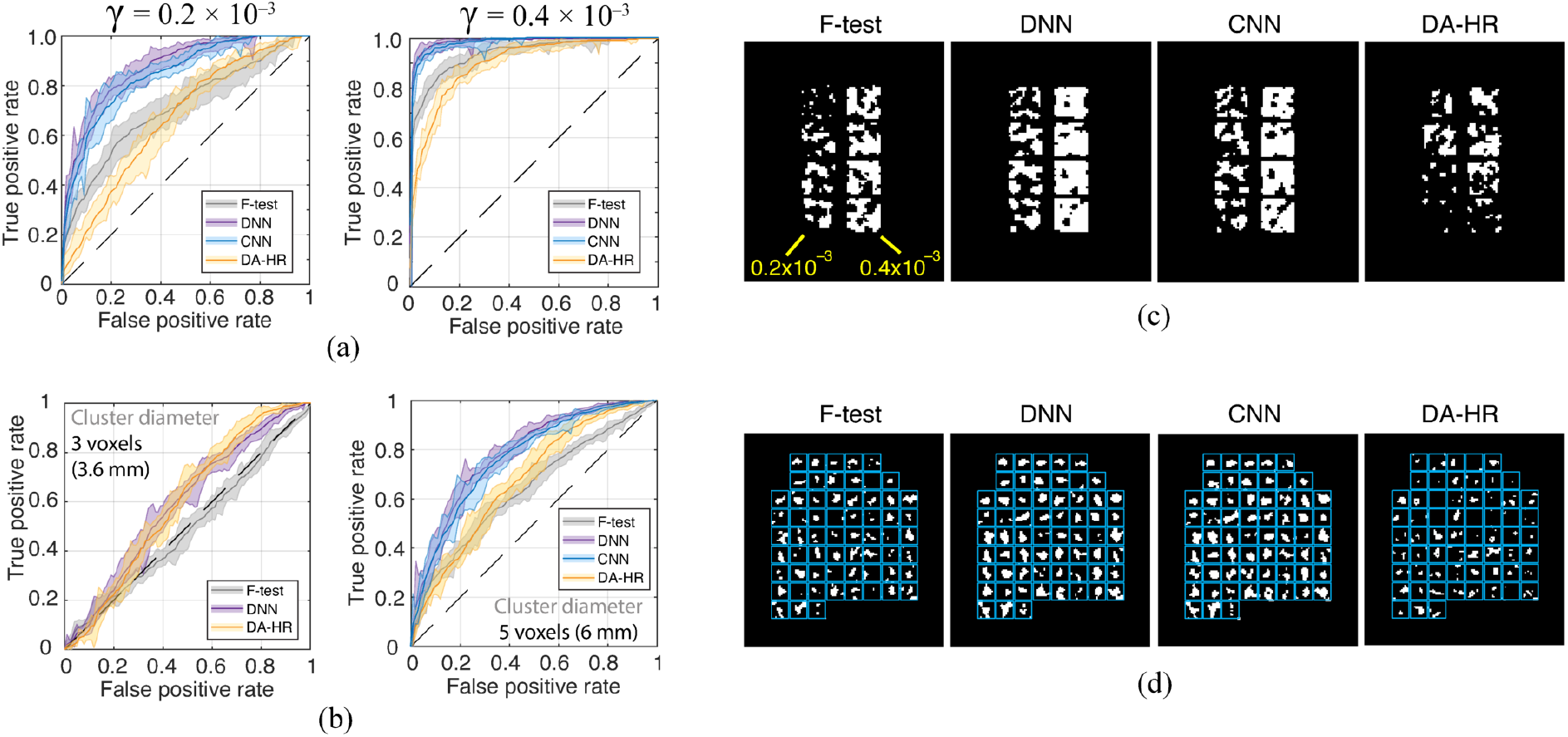
(a) Examples of ROC curves for the tested methods measured in the uniform TNR regions (BP_ND_=4). Shaded regions represent the 95% confidence intervals, computed using a bootstrap procedure with 50 trials. (b) Examples of ROC curves measured in the TNR cluster region (BP_ND_=4, *γ*=0.6× 10^−3^). Shaded regions represent the 95% confidence intervals. The CNN curve for 3-voxel clusters was omitted for clarity. (c) Detected TNR image in uniform TNR regions (FP rate 0.01). Numbers indicate *γ* values. (d) Detected TNR image in the TNR cluster region (FP rate 0.01). Rectangular cells are drawn around the locations of ground truth TNR clusters.

Examples of the ROC curves for voxels in the TNR clusters are plotted in Fig. 4b for 3- and 5-voxel clusters, and *γ* = 0.6×10^−3^. The curve shapes reveal that the ANN AUC gains over the F-test are mostly coming from the intermediate FP rates (0.2–0.8), which are unlikely to be acceptable in practical applications.

### 3.2 TNR detection sensitivity

Images showing the results of TNR detection in the uniform and cluster test regions are plotted in Fig. 4c and 4d, respectively. The images demonstrate the improved sensitivity of DNN and CNN compared to the F-test. DA-RES images (not shown) were similar to the DNN images. For the *γ* values of 0.4 × 10^−3^ and greater, most voxels in the uniform regions were classified correctly. Most of misclassified TNR voxels were near the edges, likely due to the partial volume effect (from the combined intrinsic resolution blurring and denoising). Interestingly, the locations of the false negative voxels were very similar in the F-test and ANN-produced TNR images, suggesting that there were locations in the input image that were consistently difficult to correctly classify for all methods. The detected TNR clusters (Fig. 4d) were noticeably more prominent in the DNN and CNN images compared to the F-test and DA-HR.

The TNR detection sensitivities for voxels in the uniform regions are given in Table 1, for the fixed FP rate of 0.01. The DNN and DA-RES methods offered the most substantial gains in sensitivity compared to the F-test, especially for intermediate *γ* values of 0.4× 10^−3^ and 0.6× 10^−3^. The greatest improvement was found with BP_ND_=5 and *γ*=0.4× 10^−3^: the sensitivity was 0.62 for the F-test and 0.9 for DNN/DA-RES. On the other hand, the sensitivity of DA-HR was very low, particularly for BP_ND_=3.

**Table 1:** TNR Detection Sensitivity in Uniform Regions (FP rate = 0.01)

| | $\gamma$ (s<sup>-1</sup>) | | | |
| --- | --- | --- | --- | --- |
|  | 0.2×10<sup>-3</sup> | 0.4×10<sup>-3</sup> | 0.6×10<sup>-3</sup> | 0.8×10<sup>-3</sup> |
| BP<sub>ND</sub> = 3 |  |  |  |  |
| **F-test** | 0.12 | 0.38 | 0.62 | 0.85 |
| **DNN** | 0.19 | 0.6 | 0.82 | 0.95 |
| **CNN** | 0.15 | 0.51 | 0.77 | 0.91 |
| **DA-RES** | 0.19 | 0.57 | 0.8 | 0.95 |
| **DA-HR** | 0.023 | 0.087 | 0.15 | 0.28 |
| BP<sub>ND</sub> = 4 |  |  |  |  |
| **F-test** | 0.15 | 0.6 | 0.82 | 0.97 |
| **DNN** | 0.23 | 0.81 | 0.96 | 0.99 |
| **CNN** | 0.22 | 0.78 | 0.95 | 1.0 |
| **DA-RES** | 0.22 | 0.79 | 0.95 | 0.99 |
| **DA-HR** | 0.055 | 0.35 | 0.35 | 0.53 |
| BP<sub>ND</sub> = 5 |  |  |  |  |
| **F-test** | 0.2 | 0.62 | 0.88 | 0.99 |
| **DNN** | 0.37 | 0.90 | 0.98 | 1.0 |
| **CNN** | 0.35 | 0.89 | 0.98 | 1.0 |
| **DA-RES** | 0.41 | 0.90 | 0.99 | 1.0 |
| **DA-HR** | 0.076 | 0.21 | 0.39 | 0.48 |

### 3.3 Detection of true TNR clusters

Using the test image with TNR clusters, we measured the average detected fraction of a TNR cluster as a function of the cluster size and *γ* value. The results are provided in Table 2. Only those clusters where at least 7 central voxels were detected correctly entered the calculation; this explains the artificially higher values for the 3-voxel clusters with 0.4×10^−3^. For greater *γ* values and larger clusters, we found that DNN, CNN and DA-RES detected a larger fraction than the F-test by a factor of approximately 1.7. In the best-case scenario (*γ*=0.8 × 10^−3^, cluster diameter 7 voxels), we could detect only about 50% of a TNR cluster. The Dice coefficient was measured for the detected clusters to take both the sensitivity and specificity into account, i.e. what fraction of the baseline voxels around the TNR cluster was classified correctly (Fig. 5a). For all *γ* values, methods DNN, CNN and DA-RES had greater Dice coefficients than the F-test for 5- and 7-voxel TNR clusters.

**Table 2:** Fractions of Detected TNR Clusters (FP rate = 0.01)

|  | Cluster diameter |  |  |
| --- | --- | --- | --- |
|  | 3 vox. (3.6 mm) | 5 vox. (6 mm) | 7 vox. (8.4 mm) |
| $\gamma = 0.4 \times 10^{-3}$ | | | |
| **F-test** | **0.16** ± 0.11 | **0.07** ± 0.094 | **0.11** ± 0.089 |
| DNN | **0.19** ± 0.23 | **0.075** ± 0.078 | **0.17** ± 0.13 |
| CNN | **0.16** ± 0.17 | **0.093** ± 0.085 | **0.18** ± 0.14 |
| DA-RES | **0.25** ± 0.27 | **0.082** ± 0.084 | **0.18** ± 0.13 |
| DA-HR | **0.082** ± 0.038 | **0.086** ± 0.081 | **0.089** ± 0.077 |
| $\gamma = 0.6 \times 10^{-3}$ | | | |
| **F-test** | **0.14** ± 0.083 | **0.13** ± 0.12 | **0.21** ± 0.11 |
| DNN | **0.14** ± 0.12 | **0.18** ± 0.16 | **0.34** ± 0.17 |
| CNN | **0.16** ± 0.16 | **0.18** ± 0.16 | **0.35** ± 0.16 |
| DA-RES | **0.12** ± 0.11 | **0.2** ± 0.17 | **0.34** ± 0.17 |
| DA-HR | **0.14** ± 0.11 | **0.097** ± 0.084 | **0.13** ± 0.099 |
| $\gamma = 0.8 \times 10^{-3}$ | | | |
| **F-test** | **0.17** ± 0.12 | **0.15** ± 0.12 | **0.28** ± 0.16 |
| DNN | **0.19** ± 0.13 | **0.26** ± 0.15 | **0.45** ± 0.17 |
| CNN | **0.16** ± 0.08 | **0.26** ± 0.15 | **0.47** ± 0.17 |
| DA-RES | **0.15** ± 0.11 | **0.26** ± 0.15 | **0.46** ± 0.17 |
| DA-HR | **0.16** ± 0.12 | **0.13** ± 0.12 | **0.17** ± 0.12 |

With $\gamma = 0.2 \times 10^{-3}$ , the detected cluster fraction was negligibly small with all methods.

**Figure 5:**
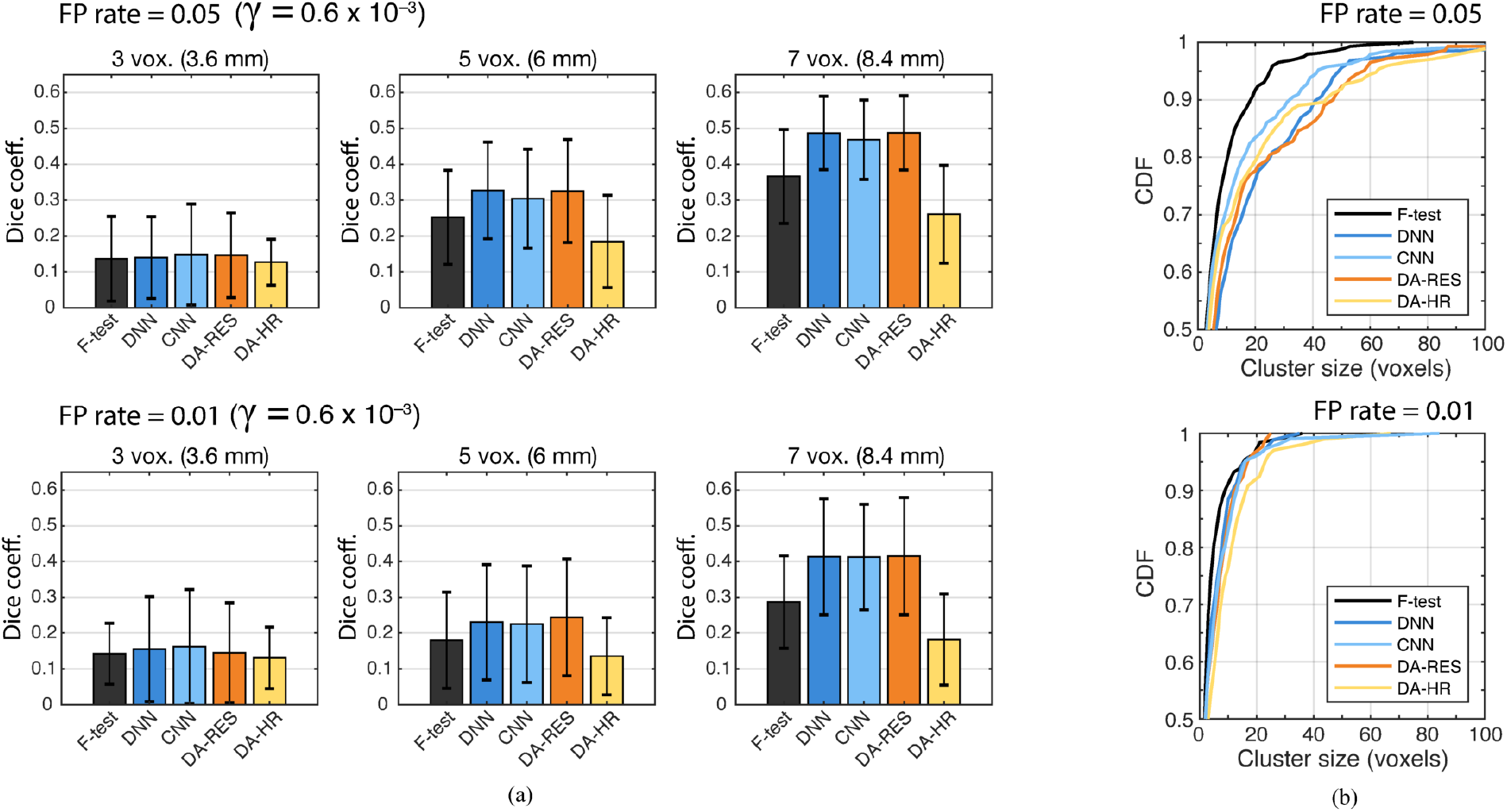
(a) Dice coefficients for the detected TNR clusters at FP rates of 0.05 and 0.01. (b) Cumulative distribution functions (CDF) of FP clusters according to their volume, at FP rates of 0.05 and 0.01 (BP_ND_=4).

### 3.4 FP cluster analysis

FP clusters were analyzed in the uniform baseline regions with BP_ND_=4, and single-voxel FP rates set to 0.05 an 0.01. Smaller FP clusters are preferable, since those can be rejected based on the cluster size thresholding [8], without sacrificing larger true positive clusters. The cumulative distribution functions of FP clusters with respect to their size are plotted in Fig. 5b. At the FP rate equal to 0.05, the 95%-tiles (99%-tiles) of the FP cluster sizes in voxels were 25 (52) for the F-test, 50 (121) for DNN, 42 (93) for CNN, 58 (106) for DA-RES, and 63 (103) for DA-HR. Thus, the ANN methods produced FP clusters that were more than twice the size compared to the F-test.

However, at the FP rate equal to 0.01, the 95%-tiles (99%-tiles) were 16 (30) for the F-test, 17 (31) for DNN, 16 (53) for CNN, 17 (24) for DA-RES, and 23 (55) for DA-HR. Thus, the FP clusters produced by the DNN and DA-RES were nearly equal in size to those produced the F-test. Thus, at the FP rate of 0.01, the methods DNN and DA-RES produced FP clusters of the same size as the F-test, while offering a substantially greater sensitivity.

### 3.5 TNR detection in the acquired image

Applying the TNR detection methods to the acquired RAC image produced parametric class-score images shown in Fig. 6a (only striatum is shown). A higher class score corresponds to a greater confidence that the voxel has non-zero TNR. Since the class-scoring functions are different in the F-test and ANN-based methods, the scores cannot be directly compared (i.e. prior to using a threshold for classification). However, the spatial distributions of the scores can be compared; for the F-test image, we log-transformed the F-scores to bring them to similar scale with the ANN class scores.

**Figure 6:**
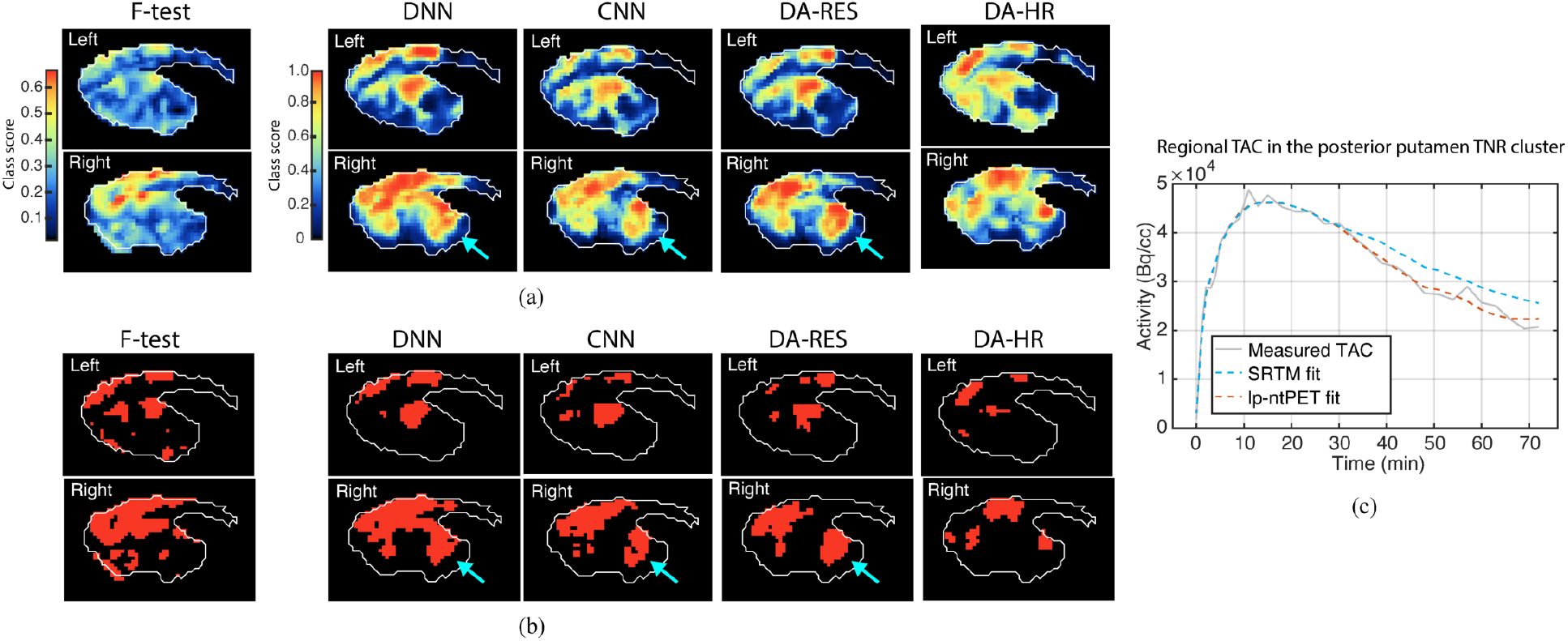
(a) Parametric class score (baseline/TNR) images for the acquired RAC scan obtained with each method. (b) Binary TNR maps obtained by thresholding the class score images. Arrows indicate the TNR cluster not detected by the F-test. (c) Measured regional TAC in the new TNR cluster detected by DA-RES, with SRTM and lp-ntPET fits.

On the left side of the striatum, the ANN methods detected similar clusters to the F-test. Namely, four high-score regions were detected, two in the superior caudate and two in the superior putamen. However, the DNN, CNN and DA-RES methods had a greater contrast of regions with high class scores as, for example, in the inferior putamen. On the right side of the striatum, DNN, CNN and DA-RES also detected similar regions to those from the F-test, however they also detected a new high-score region in the posterior putamen not present in the F-test image.

The results of thresholding the class score images to obtain the final TNR classification at FP rate of 0.01 are plotted in Fig. 6b. The detected TNR clusters in the superior caudate/putamen were similar in size between the F-test and the DNN/CNN/DA-RES methods. The newly detected TNR cluster in the posterior putamen was connected with the caudate cluster in the DNN image, and disconnected in the CNN and DA-RES images.

To test whether the additional ANN-detected cluster in the posterior putamen was a true positive or a FP, we measured the mean regional TAC in that cluster (according to DA-RES), and fitted it with SRTM and lp-ntPET (Fig. 6c). A significantly better fit was obtained with lp-ntPET, with the following fitted parameter values: *γ*=0.226×10^−3^ s^−1^, *t_D_*=31 min, *t_P_*=43.5 min, *α*=0.25, BP_ND_=4.46. These numbers suggest that there was likely transient dopamine release in the posterior putamen during the scan, which was detected by the pANNs and missed by the F-test.

## 4 Discussion

Traditionally, the detection of TNR from dynamic PET data has been achieved by statistically comparing the lp-ntPET and baseline model fits using an F-test. Our results demonstrate that pANNs, trained to detect TNR in single-voxel TACs, outperform the F-test in terms of several objective metrics. Specifically, for a given FP rate, neural networks had higher detection sensitivity in uniform TNR regions as well as TNR clusters. When tested on a real image, ANN-detected TNR clusters included those detected with the F-test. An additional TNR cluster was detected by the DNN, CNN and DA-RES networks, which was verified to likely contain TNR by direct lp-ntPET model fitting. This outcome provides an initial validation of the approach where a neural net is trained entirely on simulated data and applied to detect TNR in real data.

For all methods, the TNR detection performance for a given input function (injected activity) increased with greater *γ* and cluster size, while the effect of the BP_ND_ magnitude was relatively low. The size of the TNR was found to be by far the strongest factor impacting the detection performance. The pANNs were substantially more effective than the F-test in detecting TNR clusters. However, for large TNR regions and relatively high *γ* values the performance was similar between all methods.

We attribute the overall better pANN performance to several factors. First, as universal approximators the ANNs can accurately capture the input data distribution. In the process of training, the ANNs have presumably encapsulated the differences between lp-ntPET and SRTM models on a frame-by-frame level. In addition, the networks likely “learned” to account for the simulated resolution blurring, which may explain their better performance in detecting smaller TNR clusters. Second, better performance may be attributed to the better handling of frame-dependent noise properties. Finally, we believe that training the ANNs using a specific input function (reference TAC), what we call ANN “personalization”, contributes to their greater sensitivity.

The most common approach to train generalizable ANNs for medical image analysis is to diversify the training set, in the attempt to train the network to handle images of varying properties (different subjects, noise levels, resolutions, etc.). In contrast, our approach entails training a separate neural net for each set of dynamic images. This becomes feasible when the generated training data closely match the acquired data. The network training time is not a limiting factor, as small networks such as the ones used here can be trained on modern graphics processing units within minutes. On the other hand, physical effects such as the scattered and random coincidences can introduce a consistent bias between simulations and real scans when not fully or correctly taken into account. Another confounding factor is that real tracer kinetics may not be well-approximated by the used baseline and TNR models, however this limitation is also intrinsically present in the F-test approach. In addition, there are always unaccounted differences between the intended scan protocol and what happens in reality. The exact level of the required match between the simulated and real data still needs to be investigated in detail. In this work, the described methodology to generate simulated data yielded a relatively good generalization performance, as evidenced by the patterns in Fig. 6b (excluding the newly detected cluster).

Since the ground truth is unknown with real dynamic scans, ANN training for TNR detection will have to rely either on simulations, or ground truth proxies. With using simulated data for training, there will always be a concern regarding the generalization of measured performance to real data. However, within our approach, the simulated-to-real generalization only has to be achieved only for a specific scan, which is arguably an easier task compared to trying to generalize across different subjects, scanners and protocols. An interesting observation is the fact that by personalizing the ANNs there is no apparent need to worry about generalization and algorithm fairness, i.e. known and unknown biases related to subject age, weight, and other relevant attributes [27].

Interestingly, DNN and CNN performed rather similarly despite the different architectures. This may be a consequence of our ANNs being relatively small and shallow. Likewise, the practically identical performance of DNN and DA-RES demonstrates that there is no benefit in performing autoencoder-based TAC denoising prior to classification. The relatively worse performance of DA-HR compared to the other ANNs likely stems from data compression in the hidden representation. We speculate that within the DA bottleneck layer, a portion of TNR-related information is removed or marginalized due to the fact that the DA is trained using only the baseline data. Therefore, hidden TAC representation would have less available information, i.e. fewer salient features related to TNR compared to a network that is trained on the full input. Thus, using a hidden DA representation specifically for TNR detection may be suboptimal. However, this architecture could be attractive in situations where the primary objective is different (e.g. denoising) to detect unexpected changes in the input distribution. In a recent study not related to medical imaging, an autoencoder was trained by optimizing a modified cost function that enabled the bottleneck layer to capture anomalies more efficiently [26]; it may be of interest to adopt this technique for medical image processing and analysis.

A limitation of our method is that each time the scan or stimulus protocols change, the simulation parameters need to be adjusted accordingly. We tested one particular protocol; however, the gains in performance over the F-test may not be consistent across all possible protocols. Another limitation of our study is that we only tested the pANNs on data from one real scan. However, the objective of this work to perform a thorough method testing with simulated data, where the ground truth is known. We demonstrated that generalization from simulated to real data is possible in principle; future studies will focus on analyzing the real detected TNR in multiple subjects.

To summarize, we believe that using pANNs for TNR detection is a viable approach under the generalization caveats discussed above. In this work we only tested ANNs with one-dimensional inputs, to be able to compare them fairly to the F-test. On the other hand, training ANNs that operate on 4D data is likely to lead to further improvements in TNR detection performance. Current results can serve as a guide for the development of more advanced architectures and testing them on a larger number of real images.

## 5 Conclusion

In this work, we have tested several ANN architectures for the task of voxel-level TNR detection in simulated and real human PET imaging data, and compared the ANN performance to the standard detection method based on the F-test. Starting from a real RAC image, we trained “personalized” classifier ANNs using simulated data that have been matched with the acquired image in terms of the signal and noise. Results obtained with simulated test images demonstrate that ANNs can significantly outperform the F-test in terms of several objective classification metrics. When applied to the real image, the ANNs identified an additional TNR cluster missed by the F-test. These results demonstrate that ANNs represent a promising approach to identify voxel-level dopamine release and other types of TNR with high sensitivity.

## Acknowledgment

This work was supported by the Natural Sciences and Engineering Research Council of Canada grant number 240670-13, and Canadian Institutes of Health Research grant number 125989.

